# A toxin-antitoxin system associated transcription factor of *Caulobacter crescentus* can influence cell cycle-regulated gene expression during the SOS response

**DOI:** 10.1101/2020.07.22.216945

**Authors:** Koyel Ghosh, Kamilla Ankær Brejndal, Clare L. Kirkpatrick

## Abstract

Toxin-antitoxin (TA) systems are widespread in bacterial chromosomes but their functions remain enigmatic. Although many are transcriptionally upregulated by stress conditions, it is unclear what role they play in cellular responses to stress and to what extent the role of a given TA system homologue varies between different bacterial species. In this work we investigate the role of the DNA damage-inducible TA system HigBA of *Caulobacter crescentus* in the SOS response and discover that in addition to the toxin HigB affecting cell cycle gene expression through inhibition of the master regulator CtrA, HigBA possesses a transcription factor third component, HigC, which both auto-regulates the TA system and acts independently of it. Through HigC, the system exerts downstream effects on antibiotic (ciprofloxacin) resistance and cell cycle gene expression. HigB and HigC had inverse effects on cell cycle gene regulation, with HigB reducing and HigC increasing the expression of CtrA-dependent promoters. Neither HigBA nor HigC had any effect on formation of persister cells in response to ciprofloxacin. Rather, their role in the SOS response appears to be as transcriptional and post-transcriptional regulators of cell cycle-dependent gene expression, transmitting the status of the SOS response as a regulatory input into the cell cycle control network via CtrA.

**Importance:** Almost all bacteria respond to DNA damage by upregulating a set of genes that helps them to repair and recover from the damage, known as the SOS response. The set of genes induced during the SOS response varies between species, but frequently includes toxin-antitoxin systems. However, it is unknown what the consequence of inducing these systems is, and whether they provide any benefit to the cells. We show here that the DNA damage-induced TA system HigBA of the asymmetrically dividing bacterium *Caulobacter crescentus* affects the cell cycle regulation of this bacterium. HigBA also has a transcription factor encoded immediately downstream of it, named here HigC, which controls expression of the TA system and potentially other genes as well. Therefore, this work identifies a new role for TA systems in the DNA damage response, distinct from non-specific stress tolerance mechanisms which had been proposed previously.

## Introduction

Bacterial toxin-antitoxin (TA) systems have been the subject of intensive study since their widespread prevalence in prokaryotic chromosomes, as well as mobile genetic elements, was discovered (1). TA systems consist of two components, a toxin protein which can inhibit some aspect of central cellular metabolism and an antitoxin, either protein or RNA, which can inhibit the toxin activity or production at the post-transcriptional or post-translational level, depending on the type (2–6). The best characterised systems are those of type II where both toxin and antitoxin are small proteins which form a non-toxic complex with each other, of variable stoichiometry depending on the TA system and the organism in which it is found. This complex usually also binds the promoter of the TA system through a DNA binding domain found in the antitoxin and transcriptionally represses it (7). This repressive activity can be modulated by the abundance of the toxin protein by “conditional cooperativity,” a mechanism by which low levels of toxin protein promote TA complex binding to the TA system promoter, while high levels of toxin destabilise TA complex binding resulting in derepression of transcription (8). The mechanisms of toxins are variable but fall into two broad groups: inhibition of DNA replication by inhibiting the activity of DNA gyrase, and inhibition of translation at various levels including cleavage or modification of mRNA, tRNA or rRNA, or phosphorylation of aminoacyl-tRNA synthetases or EF-Tu (9).

However, the physiological relevance of chromosomally encoded TA systems is still an open and hotly debated question. TA systems encoded on plasmids can certainly function as plasmid maintenance systems by elimination of plasmid free cells through post-segregational killing (PSK), where the antitoxin component of the TA complexes in a plasmid-free cell is degraded and not replaced, resulting in elimination of plasmid free cells from the population because they are poisoned by excess toxin (10). It has also been noted that chromosomal TA systems are associated with cryptic prophages (11), superintegrons (12) and transposons (13) and have been suggested to promote stability and/or propagation via horizontal gene transfer of these mobile elements. Other TA systems, of types III and IV, have been shown to have roles in bacteriophage resistance (14, 15). A third highly popular hypothesis for the role of TA systems was promoting persistence to antibiotics or other stress conditions (3). However, work that purported to show that persistence in *E. coli* was mediated by activation of multiple TA systems was subsequently withdrawn, due to the discovery that the apparent persistence phenotype was due to prophage infection during mutant construction (16). Another study which showed that the type II TA system MqsRA of *E. coli* was an important mediator of the stress response (17) could not be reproduced by other groups (18). Frequently it has been assumed that the transcriptional activation of TA systems in response to various stress conditions is an indication that they are important for the corresponding stress response, and that transcriptional activation could be used as a proxy for activation of the TA system at the functional level, but it has recently been shown that during transcriptional activation of the stress-induced type II TA systems in *E. coli*, their toxins are neither released nor active (19). Hence, it is currently unclear whether type II TA systems have any involvement in persistence at all. Some type I TA systems can indeed induce persistence through their toxins which act as membrane pores and collapse the proton motive force across the membrane, thereby leading to a shutdown of cellular metabolism (20). However, other mechanisms which lead to decreased metabolic rate and slow growth can also induce persistence, independently of TA system induction or activity (21). The current key factor in induction of persistence seems to be slow growth, regardless of the cause, and no TA system has yet been reproducibly found to be necessary or sufficient for this (2).

The persistence-inducing type I TA system mentioned above (*tisAB/istR*) was seen to induce persistence in the presence of antibiotics that cause DNA damage and induce the SOS response, and the presence of type II TA systems in the SOS response regulon has also been observed in *E. coli* (22) and in *Caulobacter crescentus* (23). In these cases, the LexA repressor binds to the promoter of the TA system in addition to the antitoxin. An unusual aspect of the *Caulobacter crescentus* LexA-regulated TA system, HigBA, was that the repression provided by LexA was hierarchically superior to that of the antitoxin, allowing deletion of the antitoxin without loss of viability. Creation of a strain lacking the antitoxin allowed the identification of the targets of the mRNA-degrading toxin HigB in *Caulobacter*, which surprisingly included the essential master regulator of the cell cycle, CtrA. This TA system was not transcriptionally induced by any other stress than DNA damaging antibiotics, and appeared to have a role in resistance to these antibiotics because deletion of the *higB* toxin gene improved viability during growth on ciprofloxacin plates, in particular for the hypersensitized Δ*lexA* strain (23). However, a role for HigBA in persistence to antibiotics was not explored at that time.

The cell cycle regulator CtrA of *Caulobacter* is responsible for the asymmetric cell cycle of this bacterium, which can be seen by the division of a predivisional cell into two genetically identical but morphologically dissimilar cell types, the stalked and swarmer cells (24, 25). CtrA is a transcription factor that regulates DNA replication by binding to the origin of replication and silencing it, as well as regulating (positively and negatively) transcription of many other genes, in concert with two other transcriptional regulators MucR and SciP (26–28). CtrA is present and abundant in swarmer cells and keeps these cells in G1 phase, where they are motile but do not replicate. In contrast, it is degraded and dephosphorylated in stalked cells to permit the onset of chromosome replication and cell division (29). The ability of the toxin HigB to cleave the *ctrA* mRNA suggested that this TA system could potentially influence the cell cycle during DNA damage conditions.

In the original paper which defined the SOS regulon of *Caulobacter* (30) it was indeed seen that *higBA* was transcriptionally upregulated in a Δ*lexA* mutant. Moreover, the gene immediately 3’ to *higBA*, the uncharacterised gene CCNA_03131, was also upregulated to a similar extent, suggesting that it might be associated with *higBA* in some way. Other type II TA systems with a third component in addition to the antitoxin have been previously characterised, where the third component is a transcription factor (11, 31), an accessory antitoxin (32) or an antitoxin-stabilising chaperone (33). We therefore aimed to characterise whether this factor was involved in regulation or function of *higBA*, either in terms of the SOS response or cell cycle control via CtrA, and whether it could be considered a true third component of the system. We have also explored further whether either *higBA* or CCNA_03131 were involved in persistence and whether the cleavage of *ctrA* mRNA by HigB results in altered CtrA-dependent gene expression. Our results support a model where CCNA_03131 is a third component of the TA system and is capable of transcriptionally repressing it, but that it also acts independently of it. We find no evidence for a role of HigBA, CCNA_03131 or the LexA SOS response repressor in persistence in *Caulobacter*. Instead, the main role of the HigBA/CCNA_03131 system appears to be in regulation of cell cycle-dependent gene expression during the SOS response.

## Materials and Methods

### General growth conditions

*Caulobacter crescentus* strains were routinely grown in peptone-yeast extract (PYE) medium at 30°C and *E. coli* strains in LB at 37°C. Antibiotics were used at the following concentrations; tetracycline at 1 μg/ml for *Caulobacter* and 10 μg/ml for *E. coli*, gentamicin at 1 μg/ml for *Caulobacter* and 10 μg/ml for *E. coli* and kanamycin at 20 μg/ml (solid media) or 5 μg/ml (liquid media) for *Caulobacter* and 20 μg/ml for *E. coli*. Ciprofloxacin was prepared as 20 mg/ml stock solution in 0.1 M HCl and in all experiments involving ciprofloxacin, an appropriate volume of 0.1 M HCl was added to the control cultures or plates. Vanillate stock solution was prepared at 50 mM stock solution (adjusted to pH 8.0 with NaOH) and used at 50 or 500 μM as indicated in the figure legends.

### Strain and plasmid construction

DNA fragments for cloning were PCR-amplified with Phusion DNA polymerase (New England Biolabs) from stationary phase cultures of wild type (WT) *Caulobacter crescentus*, using PCR primers listed in Table 1. Products were purified by agarose gel electrophoresis. Cloning of the correct region was confirmed by sequencing and plasmid stocks were maintained in *E. coli* EC100 or TOP10. Plasmids used in this work are listed in Table 2. Replicating plasmids (pMT335 and plac290 derivatives) were transferred into *Caulobacter* strains by electroporation while suicide plasmids for generation of deletion mutants (pNPTS138 derivatives) were transferred by conjugation from *E. coli* S17-1 *λpir* (34). After integration of suicide vectors by recombination through one of the homologous flanking regions, secondary recombination events were induced by counterselection on PYE agar containing 3% sucrose and mutants carrying resulting in-frame deletions were screened for by PCR. Double mutants were made by introducing the Δ*higC* or Δ*higBAC* alleles into the Δ*lexA* mutant strain. Strain numbers and genotypes are listed in Table 3.

**Table 1.**
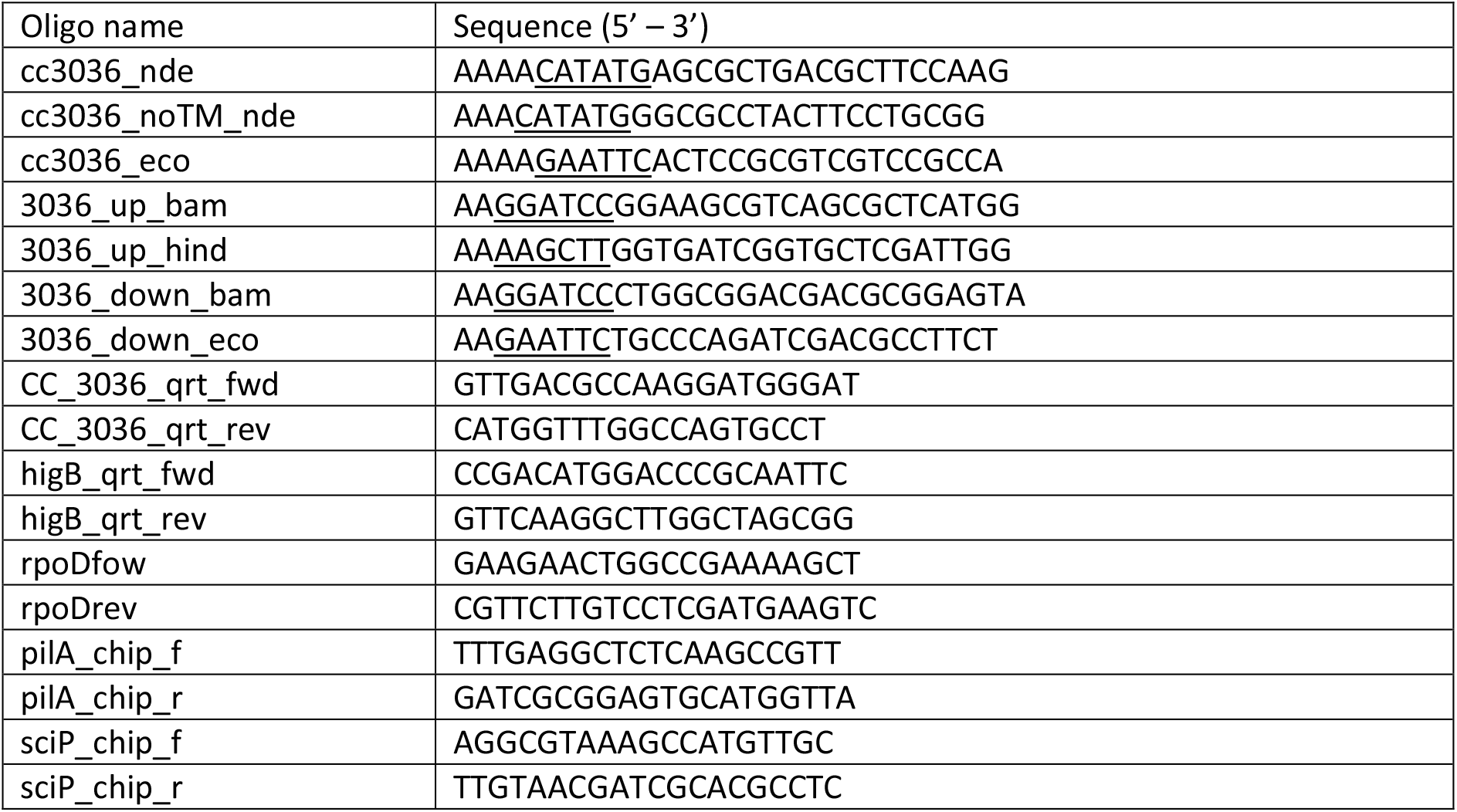
Oligonucleotide sequences used in this study. Restriction enzyme sites incorporated for cloning purposes are underlined.

**Table 2.** Plasmids used in this study.

| Plasmid name | Description | Source |
| --- | --- | --- |
| pP<sub>higBA</sub>-lac290 | plac290 (oriV, Tet<sup>R</sup>, lacZ low copy transcriptional fusion vector) with promoter of *higBA* inserted upstream of *lacZ* | (23) |
| pP<sub>pilA</sub>-lac290 | plac290 with promoter of *pilA* inserted upstream of *lacZ* | (26) |
| pP<sub>sciP</sub>-lac290 | plac290 with promoter of *sciP* inserted upstream of *lacZ* | (26) |
| pP<sub>flgB</sub>-lac290 | plac290 with promoter of *flgB* inserted upstream of *lacZ* | (26) |
| pP<sub>ctrA</sub>-lac290 | plac290 with promoter of *ctrA* inserted upstream of *lacZ* | (26) |
| pMT335 | pBBR1 ori, rep, mob, Gent<sup>R</sup>, medium copy plasmid for vanillate-inducible gene expression | (50) |
| pMT335-*higC* | pMT335 derivative, P<sub>van</sub>-*higC* | This work |
| pMT335-*higC-noTM* | pMT335 derivative, P<sub>van</sub>-*higC-noTM* (Δ1-132 amino acids, Val133Met) | This work |
| pMT335-*ctrA-DN* | pMT335 derivative, P<sub>van</sub>-*ctrA-DN* (D51E CterDD) | (23) |
| pNPTSΔ*higC* | pNPTS138 (colEI ori, M13 ori, oriT, Km<sup>R</sup>, *sacB*, suicide vector for in-frame deletions) derivative to introduce Δ*higC* allele | This work |
| pNPTSΔ*higBAC* | pNPTS138 derivative to introduce Δ*higBAC* allele | This work |

**Table 3.**
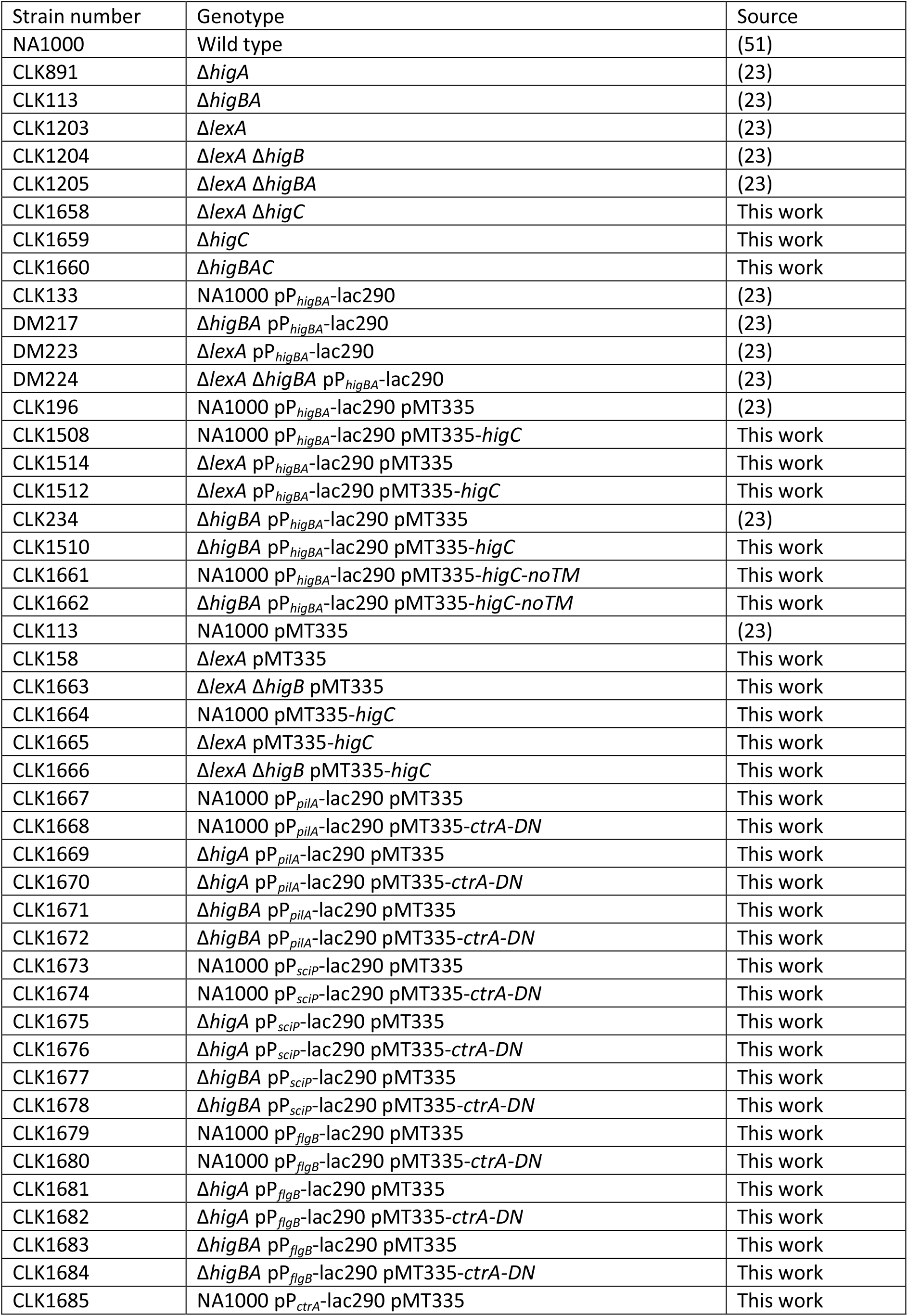
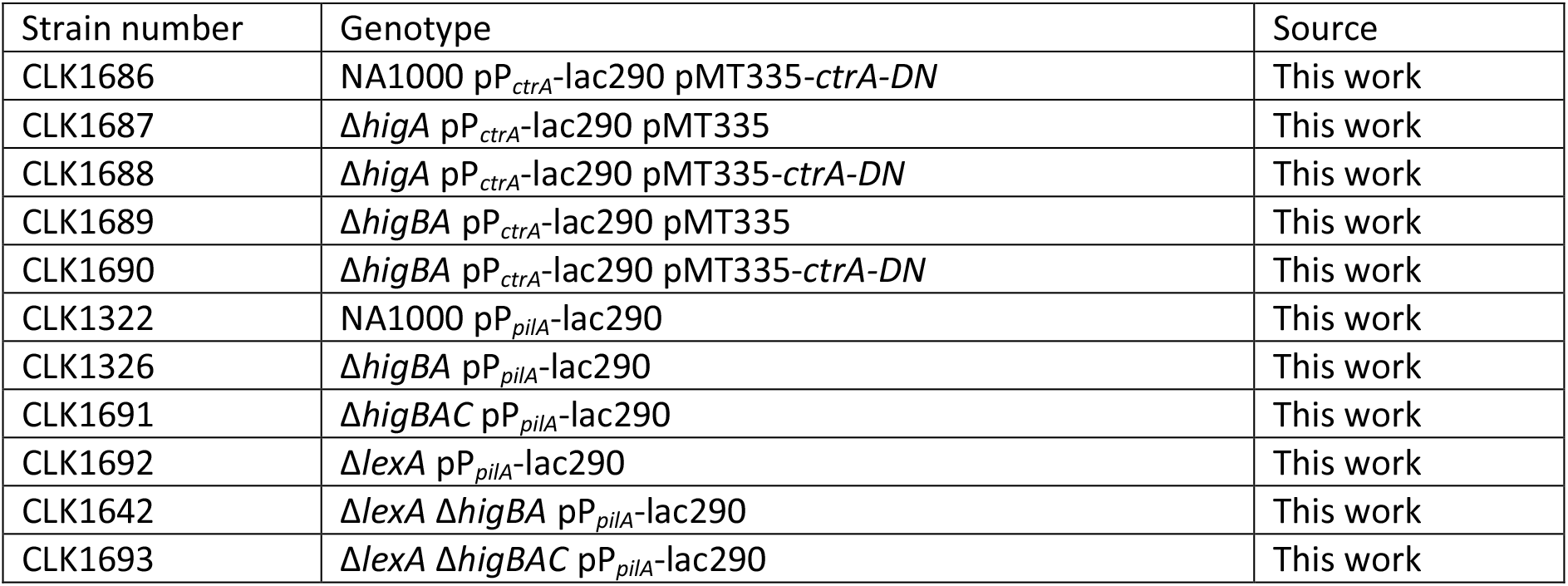
Strains used in this study.

The overexpression plasmid for WT HigC was constructed by amplification of full length *higC* with primers cc3036_nde and cc3036_eco, digested with *Nde*I and *EcoR*I and ligated into correspondingly digested pMT335 to form pMT335-*higC*. The overexpression plasmid for truncated HigC was constructed by amplification of the C-terminal half of *higC* with primers cc3036_noTM_nde and cc3036_eco. The forward primer in this reaction (cc3036_noTM_nde) contains a *Nde*I site (CATATG) which overlaps amino acid 133 (GTG-Val) and replaces it with ATG-Met. This was digested with *Nde*I and *EcoR*I and ligated into correspondingly digested pMT335 to form pMT335-*higC-noTM*. To construct the higC knockout plasmid pNPTSΔ*higC*, the flanking regions of *higC* were amplified with primer pairs 3036_up_bam and 3036_up_hind (606 bp upstream region including the first 6 amino acids of HigC) and 3036_down_bam and 3036_down_eco (556 bp downstream region including the last 6 amino acids and stop codon of HigC). These were digested with *Bam*HI/ *Hin*dIII and *Bam*HI/*Eco*RI respectively and ligated simultaneously into *Eco*RI/*Hin*dIII-digested pNPTS138. To construct the *higBAC* operon knockout plasmid, the pNPTSΔ*higBA* plasmid backbone (23) was used. This was digested with *Eco*RI/*Bam*HI to remove the fragment corresponding to the *higA* downstream region. The vector (including the upstream flanking region of *higB*) was purified by agarose gel electrophoresis and ligated to the *Eco*RI/*Bam*HI-digested PCR product of 3036_down_bam and 3036_down_eco (downstream flanking region of *higC*) to give pNPTSΔ*higBAC*.

### β-galactosidase assay

β-galactosidase assays were performed on strains carrying low copy plasmid-borne transcriptional fusions of the promoters of interest to l*acZ*. Cultures were grown to early exponential phase (OD_600_ = 0.1 – 0.4) with exposure to ciprofloxacin or vanillate as described in the main text or figure legends, followed by β-galactosidase assays using the method of Miller (35) on three independent biological replicates.

### RNA extraction and quantitative RT-PCR

RNA was extracted from 4 ml mid-exponential phase cultures which were treated with 2500 U Ready-Lyse (Bionordika) and homogenized using QiaShredder columns, prior to RNA extraction with the RNEasy Mini Kit (Qiagen) according to the manufacturers’ instructions, including on-column DNase digestion. RNA quality was assessed by agarose gel electrophoresis and concentration was measured in a Nanodrop spectrophotometer. cDNA was prepared using the SuperScript IV Reverse Transcriptase Kit (Thermo Fisher) according to the manufacturers’ instructions, on 400 ng RNA template using random hexamer primers. Quantitative RT-PCR was performed in technical and biological triplicates on a 96-well LightCycler real-time PCR system (Roche) using SYBR Green (Roche). The *higB* transcript was amplified with primers higB_qrt_fwd and higB_qrt_rev, the *higC* transcript was amplified with primers CC_3036_qrt_fwd and CC_3036_qrt_rev, and the reference gene *rpoD* was amplified with primers rpoDfow and rpoDrev. Quantification was by the standard curve method and *higB* and *higC* transcript levels were normalized to *rpoD*.

### Quantitative PCR-chromatin immunoprecipitation (qChIP)

Chromatin immunoprecipitation experiments followed by quantitative PCR were performed using an anti-CtrA polyclonal antibody (26) as described previously (36). Quantitative PCR was performed in technical duplicates on two biological replicates using primers pilA_chip_f and pilA_chip_r to amplify the promoter of *pilA* and primers sciP_chip_f and sciP_chip_r to amplify the promoter of *sciP*. Quantification was by the standard curve method and results are expressed as fold enrichment of a given product in the ChIP sample relative to the input DNA.

### Efficiency of plating assay

Resistance to ciprofloxacin and chloramphenicol was assessed by dilution spot plating. Cultures of strains to be tested were grown overnight to stationary phase, then inoculated into new medium to grow to mid-exponential phase. Culture density was measured, normalised to the OD600 of the least dense culture (OD600=0.5 or less), serially diluted in PYE to 10^−6^ and 5 μl spotted onto plates containing PYE medium with sub-inhibitory concentrations of ciprofloxacin (1 μg/ml) or chloramphenicol (0.02 μg/ml). For *higC* overexpression experiments, the plates contained gentamicin in addition to ciprofloxacin or chloramphenicol in order to maintain selection on the plasmid, and all plates contained vanillate (50 μM) to induce expression of *higC*. Plates were imaged after 3 days growth at 30°C. Images are representative of three independent biological replicates.

### Persister assay

Overnight cultures were diluted into new PYE medium and grown to mid-exponential phase (OD_600_ = 0.4 – 0.6). Ciprofloxacin was added to a final concentration of 10 μg/ml and a 100 μl sample of the culture was immediately taken out for quantification of cfu/ml at zero time. Further 100 μl samples were taken at 2, 4, 6, 24 and 48 hours after ciprofloxacin addition. Immediately after sampling, cells were washed in 1 ml PYE followed by centrifugation at 8000g for 5 minutes, repeated 3 times. Washed cells were serially diluted to 10^−6^ and plated in technical duplicates as described in the figure legends, then incubated at 30°C for 3 days. Data are reported as fraction surviving cfu/ml relative to zero time for each time point, for three independent biological replicates.

### Statistical analysis

All numerical data are reported as mean of all biological replicates performed and error bars indicate the standard deviation unless otherwise stated. Statistical significance was analysed by non-paired equal variance 2-tailed Student’s T test for comparisons between strains or treatment conditions. * signifies p < 0.05 and ** signifies p < 0.01 throughout.

## Results

### The putative transcription factor CCNA_03131 (higC) is associated with and regulates the higBA toxin-antitoxin system

We previously observed that loss of the HigB toxin in the Δ*lexA* background improved viability of the cells, likely because production or activation of this toxin is increased in the absence of LexA. Unexpectedly, we did not observe this improvement in the Δ*lexA* Δ*higBA* strain even though this also lacks HigB. Moreover, both Δ*lexA* Δ*higBA* and Δ*higBA* strains were sensitized to ciprofloxacin relative to the Δ*lexA* and wild type parent strains, respectively, while the Δ*higB* and Δ*lexA* Δ*higB* mutants were not (23). To investigate the reason for this difference, we first measured activity of a P_*higBA*_ promoter-reporter construct in wild type, Δ*lexA* and Δ*higBA* single mutants compared to Δ*lexA* Δ*higBA*. Although we observed the anticipated increased promoter activity in the absence of LexA or HigA repressors, the absence of both of these in the Δ*lexA* Δ*higBA* double mutant surprisingly led to lower, rather than higher, promoter activity (Fig 1A). Investigating the genomic context of the *higBA* TA system, we noted that a putative transcription factor gene, CCNA_03131, lies 42 bp downstream of *higBA* (Fig. 1B). Bioinformatic analysis of promoter and terminator locations (using the bprom (37) and ARNold software (38, 39), respectively) failed to find any putative promoter or terminator sequences between the 5’ end of *higB* and the 5’ end of CCNA_03131, while the same programs identified a promoter upstream of *higBA* and a putative rho-independent terminator downstream of CCNA_03131, suggesting that CCNA_03131 is a member of the *higBA* operon and potentially a third component of this TA system.

**Figure 1.**
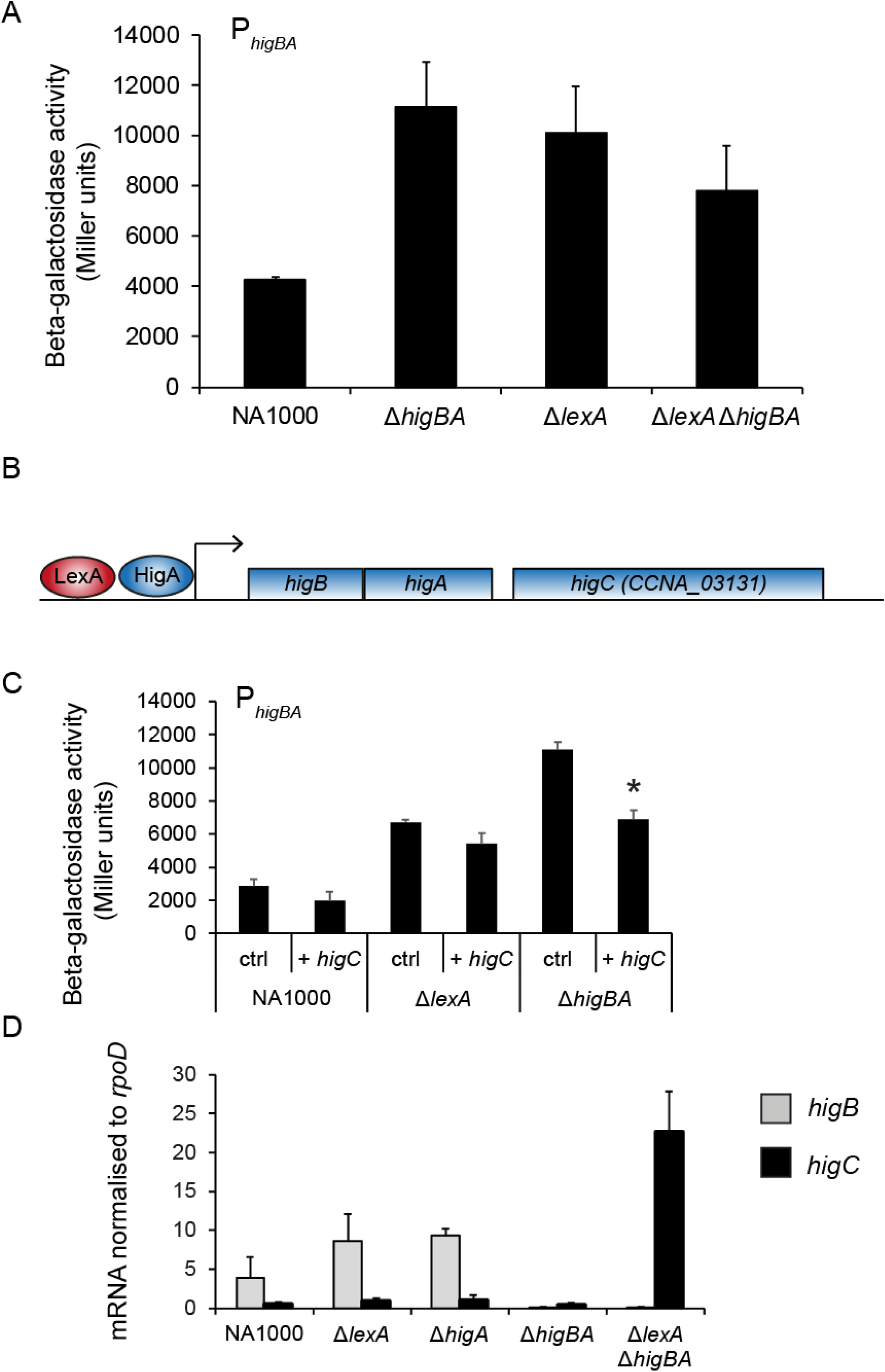
The TA system *higBA* of *Caulobacter crescentus* has a third component, *higC*. (A) Beta-galactosidase activity of P_*higBA*_-*lacZ* transcriptional reporter in NA1000 (WT), Δ*higBA*, Δ*lexA* and Δ*lexA* Δ*higBA* strains. (B) Cartoon of operon structure, approximately to scale, of *higB*, *higA* and CCNA_03101/*higC*. *higB* and *higA* are translationally coupled with a 4 bp overlap and *higC* lies 42 bp downstream of the 3’ end of *higA*. The relative positions of LexA and HigA binding at the *higBAC* promoter are based on the position of the known LexA box in the promoter (30) and on previously published ChIP-Seq data for HigA (23). (C) Beta-galactosidase activity of P_*higBA*_-*lacZ* transcriptional reporter in NA1000 (WT), Δ*lexA* and Δ*higBA* strains carrying either the pMT335 empty vector (ctrl) or the *higC* overexpression construct pMT335-*higC* (+ *higC*) after 3 hours treatment with 50 μM vanillate to induce higC expression. * indicates p < 0.05 for comparison of *higC* overexpression to empty vector control in Δ*higBA*. (D) Quantitative RT-PCR for the *higB* (grey) and *higC* (black) coding sequences performed on cDNA prepared from NA1000 (WT), Δ*higA*, Δ*higBA*, Δ*lexA* and Δ*lexA* Δ*higBA* strains. For every sample, *higB* and *higC* quantity values were normalised to the values obtained for the housekeeping gene *rpoD*. Error bars for this graph indicate standard error of the mean.

Since this gene had been annotated as a LytTR-family transcription factor (40) based on sequence homology, and some TA systems are known to have third components that act as transcription factors (11), we investigated whether it could also regulate *higBA*. Overexpressing CCNA_03131 from the vanillate-inducible promoter reduced P_*higBA*_ activity in WT, Δ*lexA* and Δ*higBA* strains relative to the empty vector control, with the most significant effect seen in the Δ*higBA* background (Fig 1C). Hence, the product of CCNA_03131 can repress the P_*higBA*_ promoter, albeit weakly, and seems to have stronger repressive activity when HigA is absent. Due to the likely co-regulation of CCNA_03131 with *higBA*, and its ability to repress transcription from the *higBA* promoter, we now consider CCNA_03131 as a part of the *higBA* TA system operon and name it *higC*. We then measured the steady-state mRNA levels for *higB* and *higC* in WT, Δ*lexA*, Δ*higA*, Δ*higBA* and Δ*lexA* Δ*higBA* strains to confirm whether their expression levels were consistent with the promoter activity measurements (Fig 1D). Both mRNAs were detectable, but *higC* appeared to be expressed at a much lower level than *higB* and no obvious induction of *higC* in the Δ*lexA*, Δ*higA*, and Δ*higBA* strains relative to WT was seen. However, *higC* was very strongly expressed in the Δ*lexA* Δ*higBA* mutant, in which the in-frame deletion of *higBA* has placed the *higC* coding sequence immediately downstream of the *higBA* promoter, and the LexA and HigA repressors are missing. Taken together, these data show that the product of *higC* functions as a repressor of the *higBAC* promoter and is strongly overproduced in the Δ*lexA* Δ*higBA* mutant, providing a plausible explanation for why the P_*higBA*_ promoter-reporter activity in this strain was unexpectedly low.

### The N-terminal helical domain of HigC is required for promoter regulatory activity

Based on sequence homology, HigC belongs to the LytTR family of DNA binding proteins (pfam04397, COG3279), but in addition to the DNA binding domain that is typical of this family, it was previously proposed to contain four transmembrane helices (40). Analysis of the HigC protein sequence by the Dense Alignment Surface (DAS) program (41) agreed with this study, suggesting that the four transmembrane helices were in the N-terminal half of the protein sequence, preceding the DNA binding domain that is predicted to start at amino acid 171 (Fig 2A). Since the existence of transmembrane helices seemed counter-intuitive in a transcription factor, which should be able to localize to the nucleoid rather than the membrane, we constructed a truncated version of HigC in which the transmembrane helices were removed (HigC-noTM). Placing this construct under the control of the vanillate-inducible promoter allowed us to compare its effect on the *higBA* promoter to wild type HigC or the empty vector control (Fig 2B). In both WT and Δ*higBA* strains, the wild type HigC repressed the promoter as before, but the truncated HigC lacking the N-terminal helical domain was completely inactive as a repressor, showing that this domain is required for activity.

**Figure 2.**
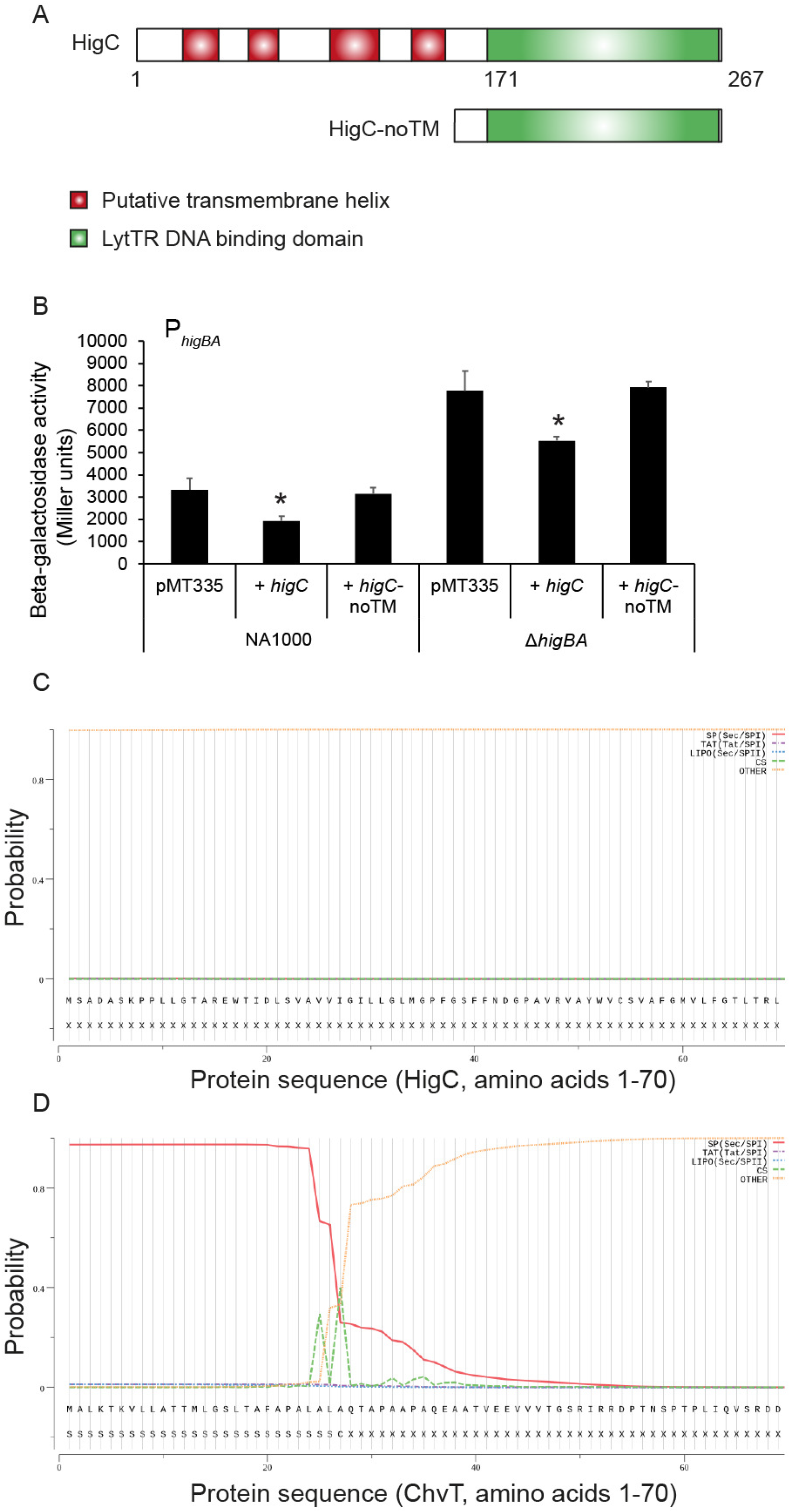
The N-terminal helical domain of HigC is needed for its activity. (A) Cartoon of HigC domain structure showing the positions of the DNA binding domain as predicted by pfam and the putative transmembrane helices as predicted by DAS, alongside the truncated variant HigC-noTM. (B) Beta-galactosidase activity of P_*higBA*_-*lacZ* transcriptional reporter in NA1000 (WT), Δ*lexA* and Δ*higBA* strains carrying the pMT335 empty vector (pMT335), the truncated *higC* overexpression construct pMT335-*higC-noTM* (+ *higC-noTM*) or the WT *higC* overexpression construct pMT335-*higC* (+ *higC*) after 3 hours treatment with 50 μM vanillate to induce higC expression. * indicates p < 0.05 for comparison of *higC* overexpression to empty vector control. (C) Signal peptide prediction using the SignalP-5.0 program set to detect signal peptides of Gram-negative bacteria, on the first 70 amino acids of HigC. The yellow line indicates that all of the provided sequence corresponds to non-signal peptide sequence with a probability of 1. (D) Signal peptide prediction using the SignalP-5.0 program as in (C) on the first 70 amino acids of ChvT. The red line indicates a high-probability Sec-dependent signal peptide and the peaks in the green line indicate two possible cleavage sites where the signal peptide could be cleaved from the rest of the protein after export.

We further analysed the HigC protein sequence for the presence or absence of a signal peptide, reasoning that if this N-terminal helical domain is genuinely a four-helix transmembrane domain, it should be preceded by a signal peptide to direct it to the membrane for co-translational insertion. However, using the SignalP software (42), no signal peptide for either the Sec or Tat secretion pathways was seen (Fig 2C). It is unlikely that this is a false negative, because the same program could detect the signal peptide of the *Caulobacter* outer membrane protein ChvT with high probability (Fig 2D). Therefore, it is possible that this domain was annotated as transmembrane helices simply because it shares the same helical secondary structure and hydrophobicity of genuine transmembrane helices, but is not actually targeted to the membrane. Since the domain was required for promoter repression activity, and bacterial DNA binding proteins frequently function as dimers or other multimers (43), we hypothesize that this domain may participate in protein-protein interactions necessary for DNA binding instead, either between HigC monomers or with other interaction partners.

### HigC affects ciprofloxacin resistance independently of the toxin HigB

We then investigated whether HigC overproduction had other phenotypic effects than P_*higBA*_ repression, by overexpressing it from the vanillate-inducible promoter in WT, Δ*lexA* and Δ*lexA* Δ*higB* strains and testing its effect on viability in the presence of antibiotics (Fig 3A). At a sub-inhibitory (for WT) concentration of ciprofloxacin, the viability of the Δ*lexA* mutant was reduced relative to the WT and Δ*lexA* Δ*higB* strains (all containing empty vector), but the viability of the three strains was unchanged on the control plate (containing gentamicin to maintain selection of the pMT335 vector) and on a sub-inhibitory concentration of chloramphenicol. However, on mild overexpression of HigC, the viability of the Δ*lexA* strain in the presence of ciprofloxacin was reduced even further, and strikingly the improved resistance of the Δ*lexA* Δ*higB* strain to ciprofloxacin was completely reversed. This effect was unique to ciprofloxacin, as it was not seen in the control condition or on chloramphenicol. Viability of a Δ*lexA* Δ*higC* strain was slightly improved relative to the Δ*lexA* parent strain on ciprofloxacin (Fig 3B), showing that the negative effect of HigC overexpression was not likely due to non-specific intolerance of producing this protein at higher levels than the cell normally experiences. Therefore, HigC negatively influences survival in the presence of DNA damaging antibiotics, especially in the context of constitutively activated SOS response of the Δ*lexA* mutant. Moreover, since this effect was observed in a Δ*lexA* Δ*higB* mutant, this effect cannot be ascribed to HigC altering P_*higBA*_ promoter activity and HigB acting as the effector of the response. Rather, HigC must be a direct effector of the ciprofloxacin sensitivity, potentially through regulatory activity on other promoters than P_*higBA*_.

**Figure 3.**
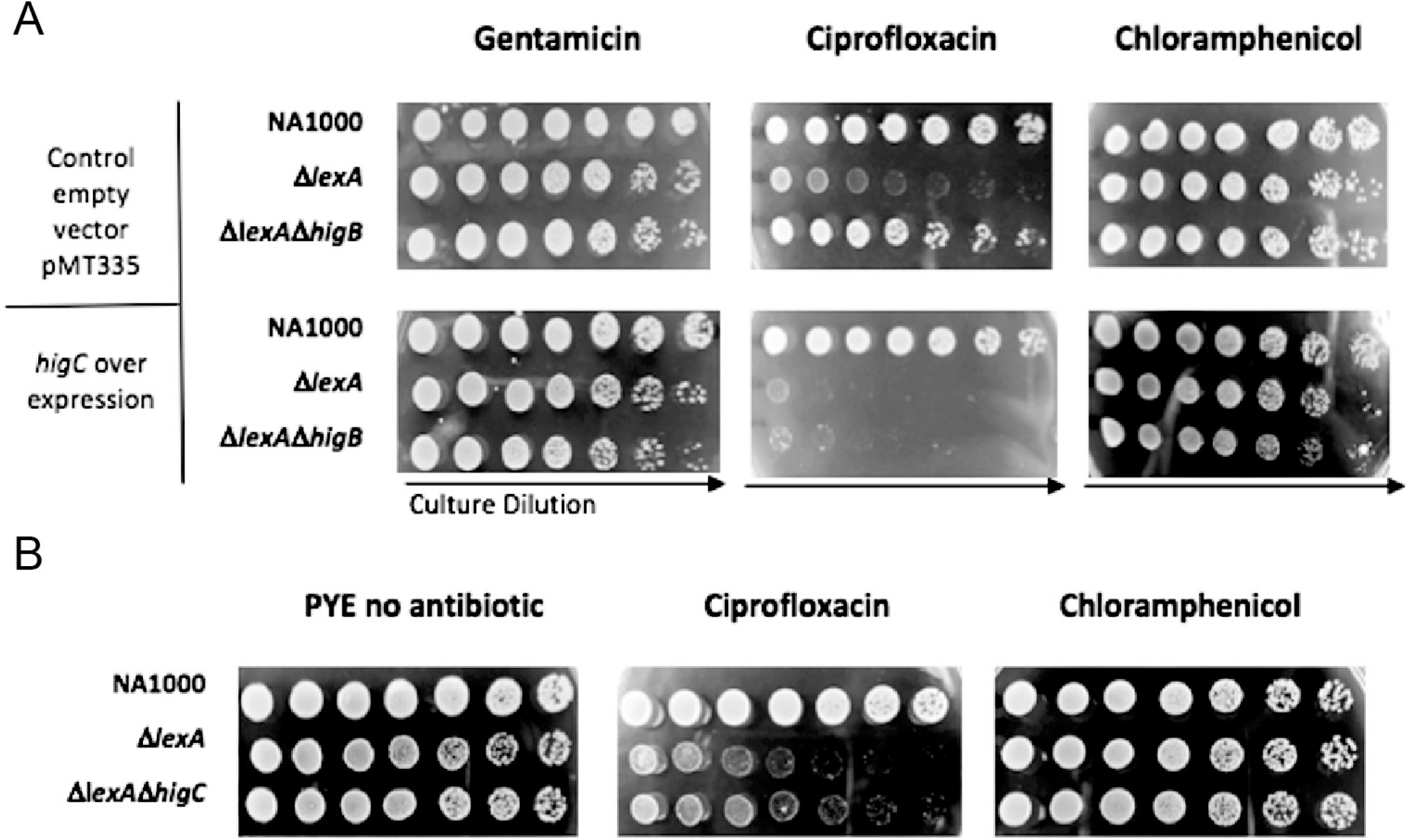
HigC influences cell viability during the SOS response. (A) Efficiency of plating assays of NA1000 (WT), Δ*lexA* and Δ*lexA* Δ*higB* strains carrying the empty vector pMT335 or the higC overexpression plasmid pMT335-*higC*, on plates containing the indicated antibiotics and 50 μM vanillate to induce *higC* expression. (B) Efficiency of plating assays of NA1000 (WT), Δ*lexA* and Δ*lexA* Δ*higC* strains. All images are representative of three independent biological replicates.

### HigBAC has no effect on formation of persister cells

Since TA systems had been previously implicated in persister cell formation, we next investigated whether the effect of HigB or HigC on viability in the presence of ciprofloxacin was associated with any change in frequency of persister cell formation. Exposure of WT, Δ*higA*, Δ*higBA*, Δ*higBAC* and Δ*higC* cells to a bactericidal concentration of ciprofloxacin followed by dilution spot plating showed that all strains exhibited a biphasic killing curve typical of persister cell formation with the initial rapid killing phase from 0 to 6 hours and with persister cells detectable after 24 and 48 hours (Fig 4A), similar to recent work in which persistence to streptomycin and vancomycin was quantified (44). This timecourse experiment showed that these strains displayed very similar biphasic curve profiles to each other with no difference in the rate of the rapid killing phase or the fraction of persisters recovered at 24 or 48 hours. However, we were unable to consistently recover persisters at 48 hours from the Δ*higBA* cultures using the spot dilution plate method, so we repeated the experiment measuring only 48-hour persisters but from a larger number of cells. This showed that all strains reproducibly had a fraction of 10^−4^ to 10^−5^ surviving persister cells after 48 hours ciprofloxacin, and that there was no significant difference in fraction of surviving persisters between any of these strains (Fig 4B). We also performed the timecourse experiment for the Δ*lexA*, Δ*lexA* Δ*higBA* and Δ*lexA* Δ*higB* strains relative to WT and found that these strains had similar biphasic curve kinetics and a similar fraction of surviving persister cells at 24 and 48 hours, and again no significant difference between any of the strains was seen (Supplementary Fig S1). Therefore, while *Caulobacter crescentus* is capable of forming persister cells upon bactericidal antibiotic (ciprofloxacin) treatment, this process is not influenced by the toxin HigB, the transcription factor HigC, or the LexA repressor which controls their expression, and the viability differences observed in our efficiency of plating assays are unrelated to persistence.

**Figure 4.**
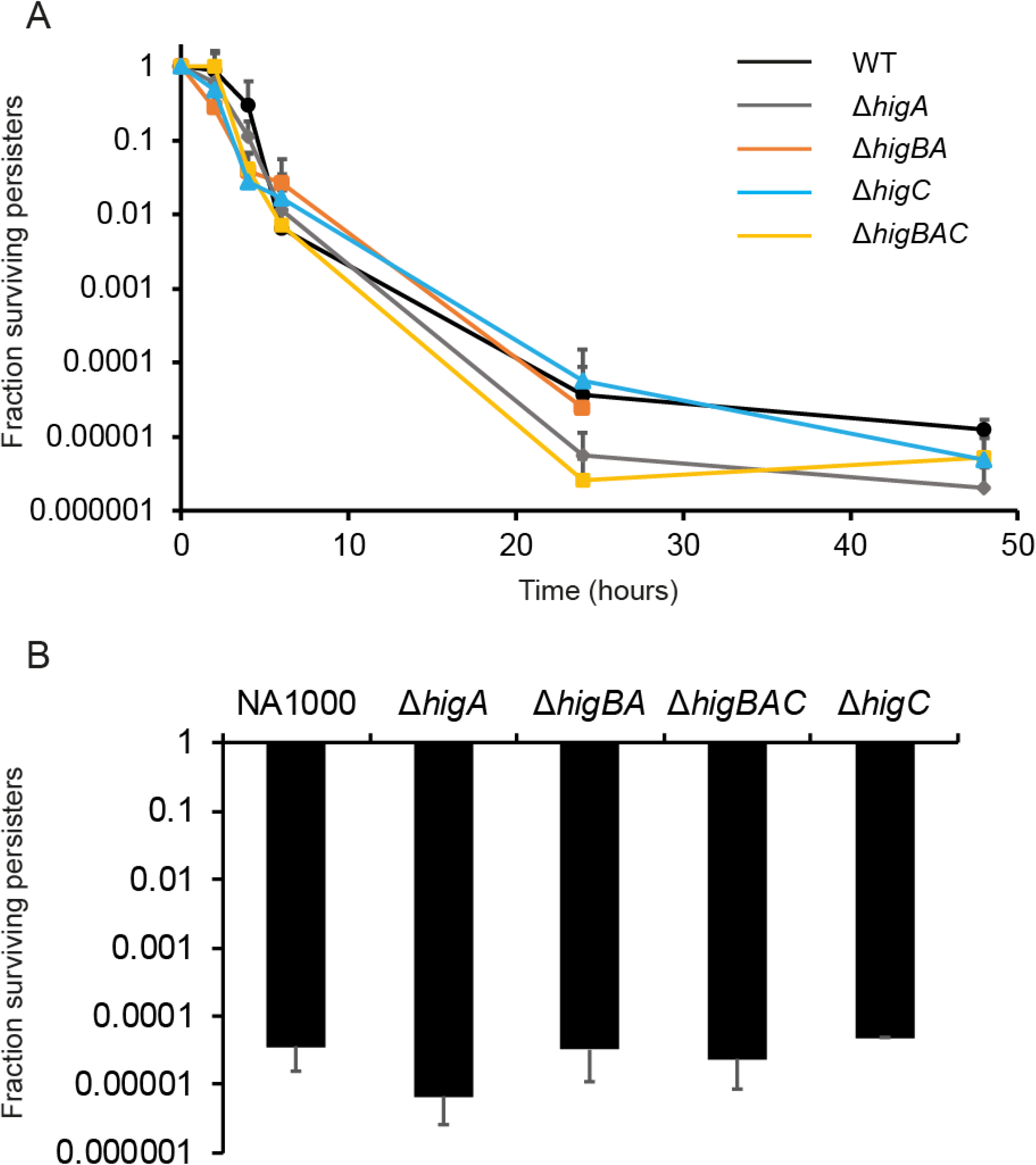
HigBAC does not influence persister cell formation. (A) Biphasic killing curve of NA1000 (WT), Δ*higA*, Δ*higBA*, Δ*higBAC* and Δ*higC* strains treated with 10 μg/ml ciprofloxacin. Samples were taken over time and spot plated (5 μl) in technical duplicates after washing and dilution to calculate surviving cfu/ml. Data were expressed as fraction surviving persisters normalised to cfu/ml at zero time, from three independent biological replicates. (B) Measurement of surviving persisters in the same strains as in (A) after 48 hr treatment with 10 μg/ml ciprofloxacin. Cfu/ml values were calculated from plating 100 μl samples of 10^−5^ and 10^−6^ dilutions of the zero-time sample and 10^0^ and 10^−1^ dilutions of the 48 hr ciprofloxacin treated sample and normalised to fraction surviving persisters at 48 hr relative to zero time.

### HigB negatively regulates CtrA-dependent gene expression

We had previously identified the transcript of the cell cycle regulator *ctrA* as a target of the toxin HigB’s mRNA interferase activity (23), and confirmed that increased HigB activity in the Δ*higA* mutant strain was associated both with a decreased proportion of swarmer cells in the population, and with protection against cell cycle arrest caused by overexpression of a dominant-negative CtrA allele (CtrA-DN) that could not be removed from the cells by regulated proteolysis (45), presumably by increased HigB-mediated degradation of the *ctrA-DN* mRNA. To investigate whether this was reflected at the phenotypic level by altered transcription of CtrA-dependent genes, we carried out β-galactosidase assays of CtrA-dependent promoter-reporters in WT, Δ*higA* and Δ*higBA* strains, with and without overexpression of CtrA-DN from the vanillate-inducible promoter. Since this allele forces the cells to arrest in the G1 phase, promoter activity in the presence of the empty vector indicates that of the mixed population, while promoter activity in CtrA-DN-overexpressing cells indicates the level of activity seen specifically in the swarmer cells.

We compared the activity of CtrA-dependent promoters that are expressed in the G1 phase (swarmer cells) and subject to repression by the co-repressors MucR1/2, with CtrA-dependent promoters that are expressed in the late S-phase and in G2 (stalked and pre-divisional cells) and subject to repression by the regulatory protein SciP, which includes the promoter of CtrA itself. In the CtrA-DN-overexpressing but otherwise WT cells, we anticipated that SciP protein levels should be high and that the CtrA-SciP-dependent promoters should be inactive or weakly active compared to the mixed population/empty vector control. Meanwhile, the CtrA-MucR-dependent promoters should be active in both conditions (possibly increased upon CtrA-DN overexpression). Then, any further differences in promoter activity in the Δ*higA* or Δ*higBA* backgrounds relative to WT should be accounted for by increased or decreased activity of the toxin HigB against *ctrA* mRNA. Consistent with our previous result that the loss of HigA had a reduced swarmer cell fraction in a mixed population but no difference in other cell types (23), as observed by FACS, we found reduced activity of the MucR-dependent G1-phase promoter P_*pilA*_ in Δ*higA* relative to WT both with and without CtrA-DN overexpression (Fig 5A). We did not observe the same effect for P_*sciP*_, suggesting that promoters controlling structural genes are better proxies for this effect than promoters controlling regulatory factors.

**Figure 5.**
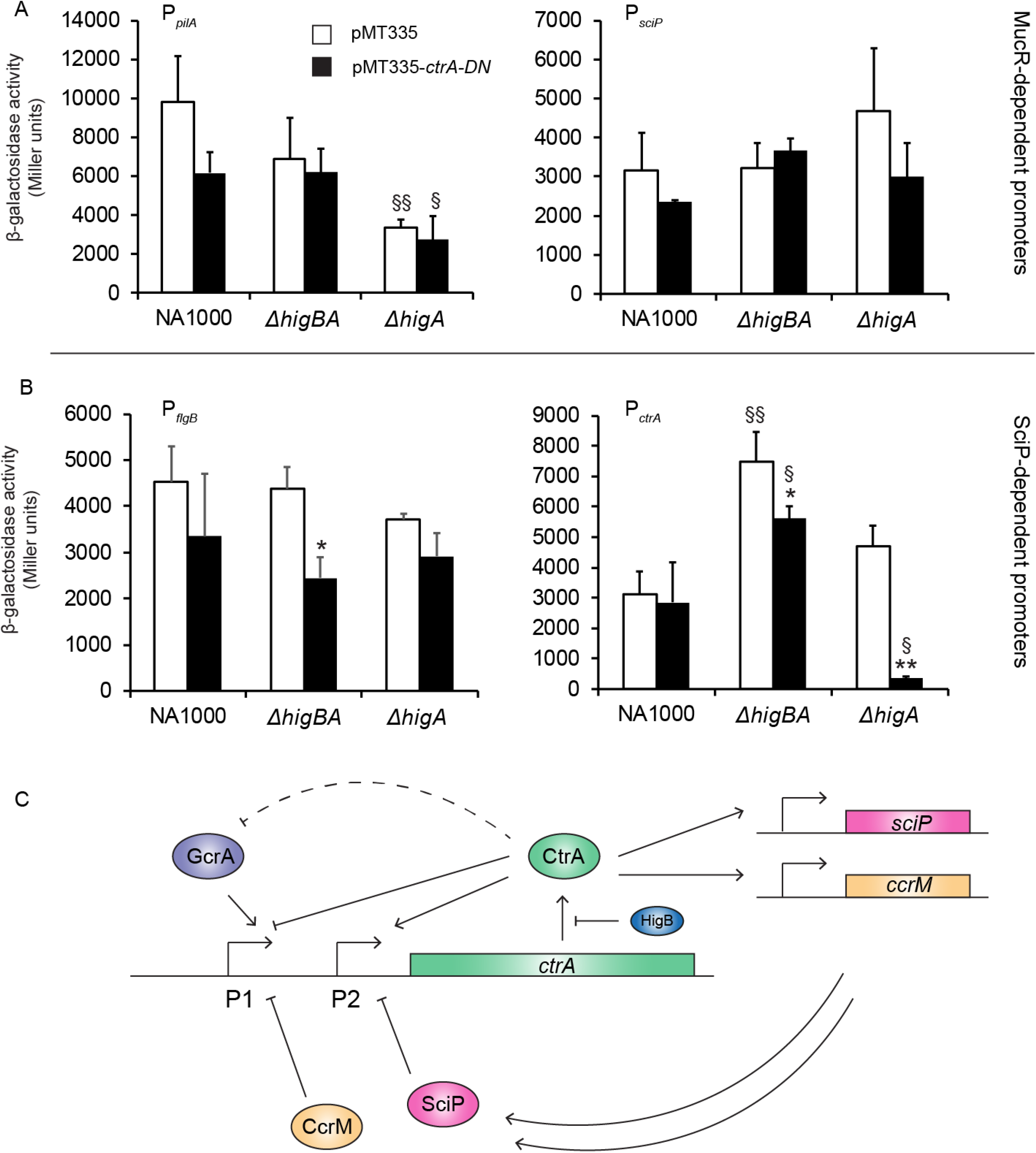
HigB activity affects expression of CtrA-dependent cell cycle genes. (A) Beta-galactosidase activity of the CtrA/MucR-dependent promoters P_*pilA*_ and P_*sciP*_ in NA1000 (WT), Δ*higBA* and Δ*higA* strains carrying either the empty vector pMT335 (white) or the non-proteolysable CtrA overexpression plasmid pMT335-*ctrA-DN* (black), after 3 hours treatment with 500 μM vanillate to induce CtrA-DN to a sufficient level to induce cell cycle arrest in the G1 phase (23). (B) Beta-galactosidase activity of the CtrA/SciP-dependent promoters P_*flgB*_ and P_*ctrA*_ in NA1000 (WT), Δ*higBA* and Δ*higA* strains carrying either the empty vector pMT335 (white) or the non-proteolysable CtrA overexpression plasmid pMT335-*ctrA-DN* (black), after 3 hours treatment with 500 μM vanillate as in (A). * and ** indicate p < 0.05 and p < 0.01 respectively for within-strain comparisons of empty vector control to *ctrA-DN* overexpression. § and §§ indicate p < 0.05 and p < 0.01 respectively for between-strain comparisons of Δ*higA* or Δ*higBA* to WT cells carrying the same plasmid. (C) Cartoon of the *ctrA* promoter and factors that regulate it. The dotted line indicates transcriptional repression of the *gcrA* promoter by CtrA, while solid lines indicate direct activation or repression.

Surprisingly, the CtrA/SciP-dependent S/G2-phase promoters were not significantly repressed in the WT background when the CtrA-DN allele was overexpressed. However, loss of HigBA appeared to promote this repression, since the Δ*higBA* strain had significantly lower activity of both promoters during CtrA-DN overexpression compared to empty vector. Surprisingly, upon CtrA-DN overexpression in the Δ*higA* strain, we observed much stronger repression of P_*ctrA*_ than in Δ*higBA* or WT (Fig 5B). The bipartite *ctrA* promoter is subject to complex multi-level regulation by SciP, CtrA itself, the S-phase associated transcription factor GcrA and the methylation state of the promoter DNA (Fig 5C), so the contribution of the multiple regulatory inputs cannot be inferred from the promoter activity measurement alone. However, we can nonetheless conclude that the Δ*higA* genetic background influences cell cycle dependent gene expression, in a manner which is consistent with its cognate toxin HigB negatively regulating *ctrA* at the post-transcriptional level. In support of this function for HigB, we also found by anti-CtrA ChIP followed by quantitative PCR that the CtrA/MucR-dependent promoters P_*pilA*_ and P_*sciP*_ had much less CtrA bound to them in non-synchronized populations of the Δ*higA* strain compared to WT and Δ*higBA* (Fig 6), despite the modest effects observed at the level of promoter activity (Fig 5A). HigB is therefore capable of negatively influencing CtrA binding to and activating its target promoters, regardless of the cell cycle phase they are associated with.

**Figure 6.**
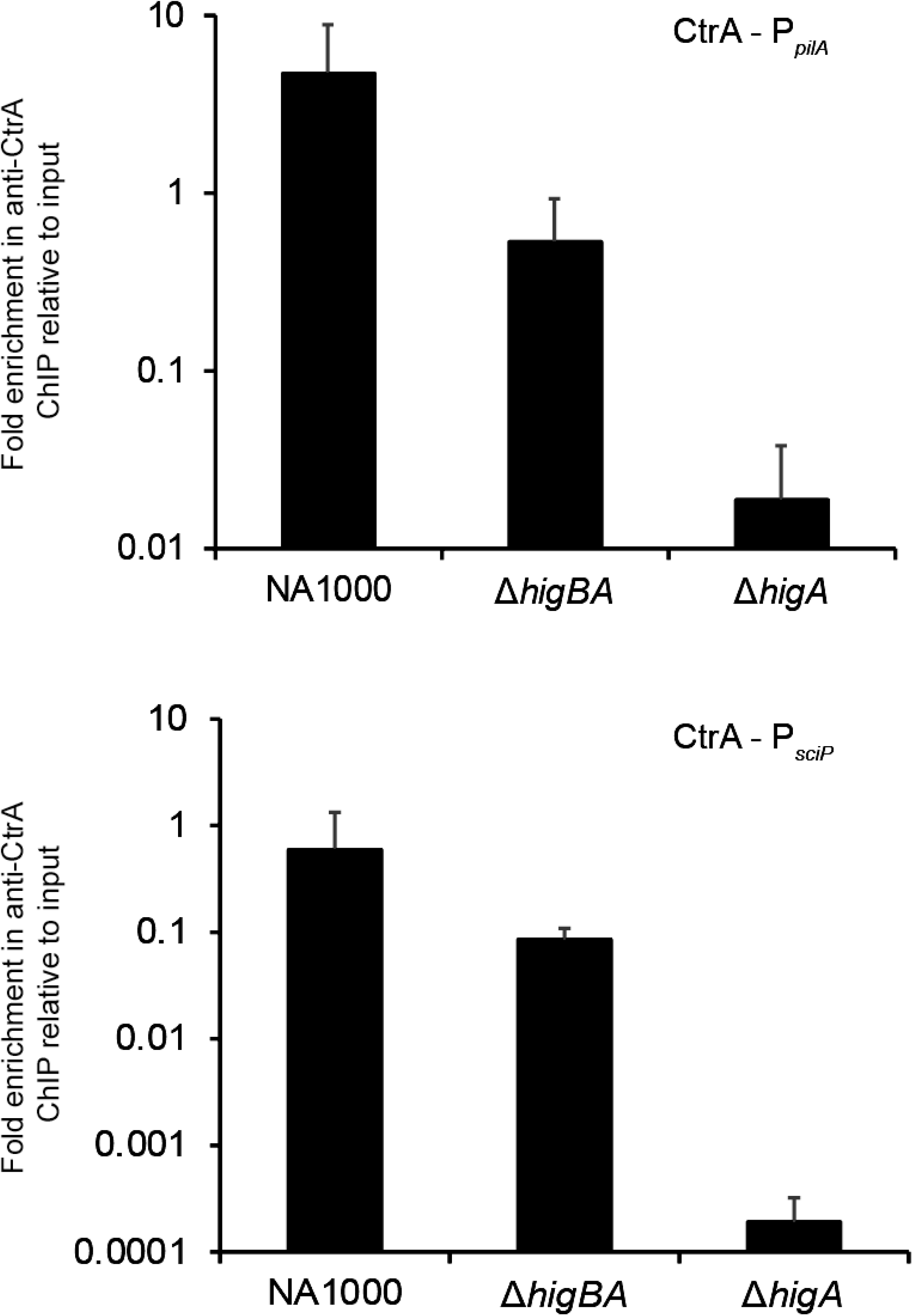
Increased HigB activity reduces binding of CtrA to its target promoters. Binding of CtrA to its target promoters P_*pilA*_ (A) and P_*sciP*_ (B) measured by ChIP with anti-CtrA antibodies followed by quantitative PCR and expressed as fold enrichment in anti-CtrA ChIP over input DNA. Data are expressed as average and standard deviation of two biological replicates, with technical duplicates performed in each experiment.

### HigC influences cell cycle gene expression during the SOS response independently of HigB

Since we had observed that HigC could negatively affect survival in the presence of DNA damaging antibiotics in a HigB-independent manner, we then investigated whether this was associated with cell cycle gene expression by using the P_*pilA*_-lacZ construct as a reporter for CtrA-dependent promoter activity in the presence and absence of ciprofloxacin, in strains lacking *higBA*, *higC* or *lexA* separately or together (Fig 7A). There was no difference in P_*pilA*_ activity between WT and Δ*higBA* in the control condition, but its activity was increased in Δ*higBA* cells treated with ciprofloxacin. However, this effect was not due to increased HigB activity in ciprofloxacin-treated WT, because the activity in a Δ*higBAC* mutant strain treated with ciprofloxacin was reduced down to WT levels again. Hence, HigC must have been responsible for the elevated P_*pilA*_ activity in the Δ*higBA* mutant upon induction of the DNA damage response with ciprofloxacin. In the Δ*lexA* strain, which has the DNA damage response constitutively activated, we observed similar results. Here, the baseline activity of P_*pilA*_ was lower, probably because of the LexA-induced cell division block (46, 47) that would prevent normal progression through the cell cycle and the associated pulse of *pilA* transcription in G1 phase. We did not observe any ciprofloxacin-induced increase in P_*pilA*_ activity in a Δ*lexA* Δ*higBA* mutant compared to the Δ*lexA* strain. However, the Δ*lexA* Δ*higBAC* quadruple mutant had decreased activity of this promoter compared to Δ*lexA* Δ*higBA*, both with and without ciprofloxacin. Therefore, in conditions where the SOS response is induced but the HigBA TA system inactive, HigC can promote expression of this CtrA-dependent promoter. This activity must be functionally independent of the HigBA TA system, in the sense that it is not mediated by HigB toxin activity against CtrA via HigC regulation of the *higBAC* promoter. Overexpression of HigC in WT cells from the vanillate-inducible promoter, under the same conditions in which we saw HigC repression of P_*higBA*_, did not result in any alteration of P_*pilA*_ activity relative to the empty vector (Supplementary Fig S2), suggesting that the effect of HigC on this promoter is either indirect, or undetectable if HigBA is present.

**Figure 7.**
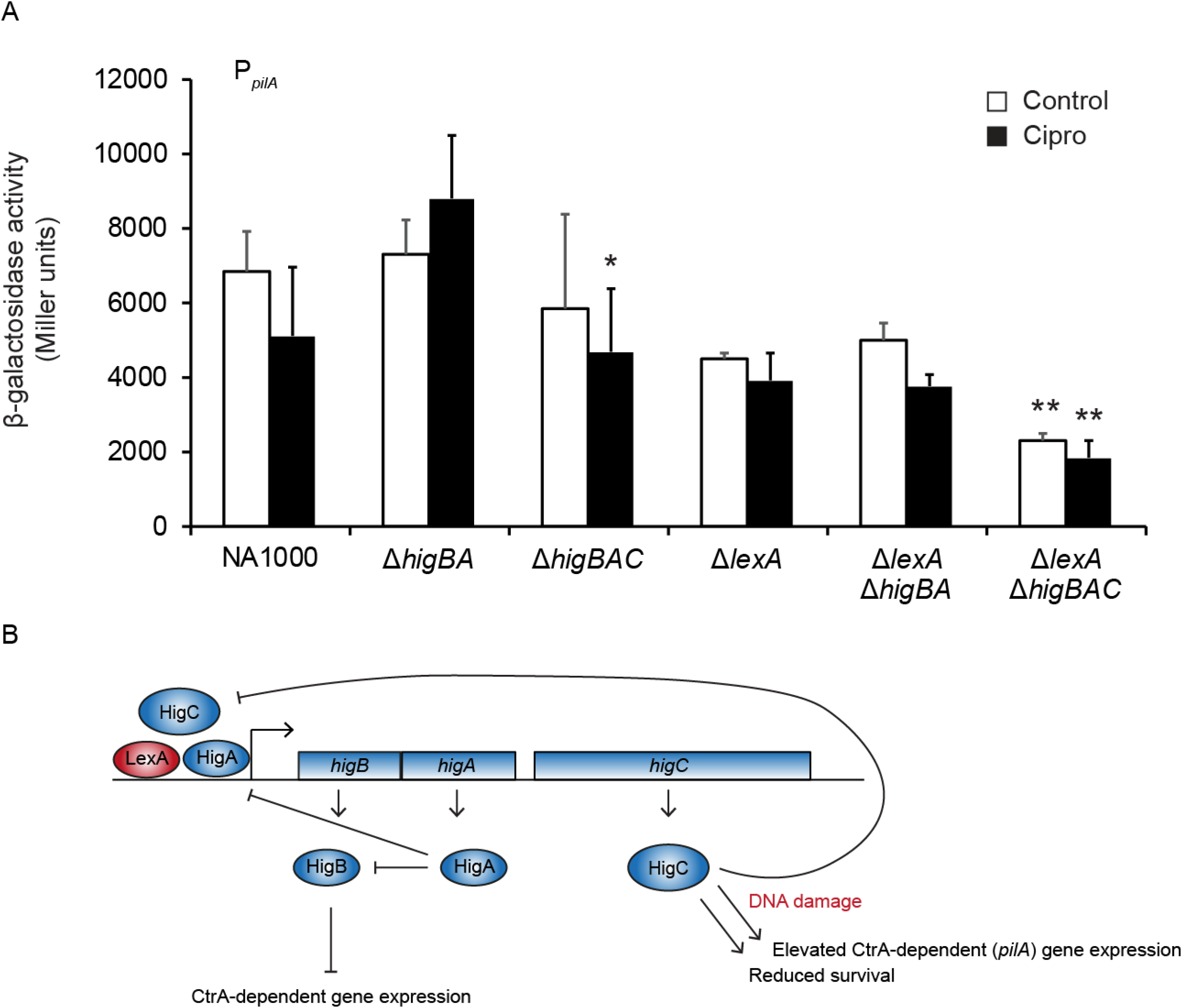
HigC affects cell cycle-regulated gene expression independently of HigB, but only during induction of the SOS response. (A) Beta-galactosidase activity of P_*pilA*_-*lacZ* in NA1000 (WT), Δ*higBA*, Δ*higBAC*, Δ*lexA*, Δ*lexA* Δ*higBA* and Δ*lexA* Δ*higBAC*, treated with vehicle (white) or with 5 μg/ml ciprofloxacin (black) for 2 hours. * indicates p < 0.05 and ** indicates p < 0.01 for between-strain comparisons of isogenic Δ*higBAC* with Δ*higBA* strains under identical treatment conditions (ciprofloxacin or vehicle). (B) Graphical summary of the regulation and function of the HigBAC 3-component TA system.

## Discussion

In the present study we report that the HigBA toxin-antitoxin system of *Caulobacter* possesses a transcription factor third component HigC, which participates in auto-regulation of the *higBA* promoter but which also acts independently of HigBA (Fig 7B). We confirm our findings from previous work that the HigB toxin can target the cell cycle regulator CtrA at the post-transcriptional level, resulting in decreased CtrA-promoter binding and lower CtrA-dependent promoter activity. Moreover, we find that under conditions of SOS response induction, HigC decreases cell viability and can also influence expression of CtrA target genes. The decrease in viability was independent of the *higB* toxin gene, while the expression of the CtrA-dependent *pilA* promoter was increased by HigC specifically in the absence of HigBA. Deletion of neither *higBA* nor *higC* had any effect on formation of persister cells in the presence of ciprofloxacin, suggesting that the HigBA- and HigC-dependent phenotypes that we observe are unrelated to the persistence phenomenon and instead indicate that HigBAC is acting as a regulatory coupling factor linking regulation of cell cycle genes to the SOS response.

While the close proximity of *higC* to *higBA* initially suggested that these genes may be in the same operon and therefore co-regulated, some aspects of whether *higC* is regulated identically to *higBA* still remain unknown. Mindful of the recent observation that in the *E. coli mqsRA* TA system, the antitoxin *mqsA* is transcribed from promoters internal to the *mqsR* coding sequence (18), we searched for promoters not only in the *higA-higC* intergenic region but in the entire *higBA* coding sequence. This bioinformatic analysis did not uncover any cryptic internal promoters. However, *Caulobacter* −10 and −35 promoter sequences do not closely match the canonical −10 and −35 boxes characterised in *E. coli* and other Gram negative bacteria, so it is also possible that this is a false negative. Indeed, in a previous global analysis of genome-wide transcription start sites over the cell cycle (48), it was found that there was a low-frequency transcription start site which corresponded to the A of the start codon of *higC*, in addition to the high-frequency transcription start site 4 bp upstream of the start codon of *higB*. Therefore, it is possible to infer that *higBA* and *higC* are transcribed from different promoters, with the *higC* promoter being much weaker, which would explain our result that the *higC* transcript is apparently present at much lower levels than *higB* in WT, Δ*higA* or Δ*lexA* strains (Fig 1D). This is more difficult to reconcile with the similar fold changes (between Δ*lexA* mutant and WT) for all three genes observed by qRT-PCR by da Rocha *et al* (30), since the region upstream of the putative *higC* transcription start site has no LexA binding site. However, if transcriptional readthrough occurred during the high levels of transcription from the *higB* promoter that would be expected during the SOS response, this could account for the *lexA*-dependent *higC* induction. Interestingly, the *higC* transcription start site was suggested to be cell cycle regulated while the *higB* transcription start site was not, with RNA-Seq reads corresponding to the *higC* site peaking at 80 to 100 minutes after synchronisation (48). This correlates closely with the peak time of the transcription start site of the *ctrA* P2 promoter, but *higC* is unlikely to be a candidate for direct regulation by CtrA since there is no CtrA binding motif (TTAA-N_7_-TTAA) in the *higA – higC* intergenic region and it has not been identified as a CtrA target in any genome-wide analysis (26, 49).

The role of HigC in repressing the HigBAC promoter is consistent with that observed for other three-component type II TA systems encoding a transcription factor (11, 31), but we also observe some unique differences. In those studies, the transcription factor was primarily responsible for repression of the system, either alone or together with the cognate TA complex acting as co-repressor. Meanwhile, for HigBAC, the HigC repression appears much less important than the repression provided by LexA and HigA. Based on our β-galactosidase data, HigC seems to exert the strongest repressive effect when HigA is absent, suggesting that it might act as a negative feedback mechanism to bring *higBAC* transcription back under control during the late SOS response, if LexA and/or HigA have been absent from the promoter. It will be intriguing to investigate how the 4-helix N-terminal domain of HigC is involved in promoter regulation, since removal of this domain completely abolished its activity as repressor of the *higBA* promoter. While we find that it is unlikely to be a true membrane protein, on account of the lack of signal peptide, one possibility is that it could mediate protein-protein interactions between HigC monomers or between HigC and other proteins. Pull-down assays of WT and truncated HigC could identify binding partners of this protein and differentiate between ones that depend on the presence of the 4-helix domain and ones that do not. Moreover, since these helical domains were identified in proteins of this family from other alpha-proteobacteria, not only *Caulobacter* (40), this domain may represent a novel conserved mediator of DNA binding protein interaction in this class of bacteria. It will also be important to define the regulon, either direct or indirect, of HigC in order to fully characterise the role of HigC in the SOS response based on which other genes it regulates in addition to *higBA*.

Our genetic approach, in which we have characterised the effect of HigB in the absence of the antitoxin, and HigC in the absence of the HigBA TA system, has allowed us to gain valuable insight into the functions of these two proteins. However, it is also important not to infer too much from studies of mutant strains about the physiological roles of these proteins in wild type cells. A criticism which is often levelled at studies of TA systems is that phenotypes of antitoxin mutant strains are not equivalent to phenotypes of wild type cells experiencing high levels of toxin production and therefore not physiologically relevant (2, 19, 21), and therefore a phenotype associated with a given TA system should only be postulated if a phenotype can be observed for a toxin or whole TA system mutant. We do indeed observe such a phenotype for HigBA, since the loss of the toxin in the Δ*lexA* background substantially improved its resistance to ciprofloxacin. However, in this work we have also found that the difference in this ciprofloxacin resistance phenotype between our Δ*higB* and Δ*higBA* strains was due to the polar effect of the *higBA* deletion on *higC* (specifically, placing it immediately downstream of the strong *higBA* promoter leading to much stronger *higC* expression than would normally occur). This underscores the importance of taking genetic context into account and not assuming that in-frame deletions are free of polar effects. Nonetheless, we can still conclude that the HigB toxin is likely to be active to some extent during the SOS response, based on the ciprofloxacin resistance phenotype of the Δ*lexA* Δ*higB* strain, and that when active it should inhibit CtrA at the post-transcriptional level resulting in lower expression levels of CtrA-activated genes. In addition, the effect of HigC overexpression or deletion on ciprofloxacin resistance of the Δ*lexA* strain shows that it can exert its effect when expressed at relatively low levels and when *higBA* is still present. Taken together, our data show that the TA system HigBAC of *Caulobacter crescentus* is a uniquely acting three-component TA system in which the toxin HigB and the transcription factor HigC exert SOS-responsive gene regulation activities at transcriptional and post-transcriptional levels, on genes involved in the cell cycle regulatory network.

## Acknowledgements

This work was supported by intramural funding from the University of Southern Denmark to CLK. We thank Patrick Viollier for the gift of the CtrA-dependent transcriptional reporter plasmids, Lykke Haastrup Hansen for technical assistance and Sergi Torres Puig for critical reading of the manuscript.

## Author contributions

KG, KAB and CLK performed experiments. KG and CLK wrote the paper. CLK conceived and designed the study.

## References

1. Pandey DP, Gerdes K. 2005. Toxin-antitoxin loci are highly abundant in free-living but lost from host-associated prokaryotes. Nucleic Acids Res 33:966–76.

2. Fraikin N, Goormaghtigh F, Van Melderen L. 2020. Type II Toxin-Antitoxin Systems: Evolution and Revolutions. Journal of Bacteriology 202:e00763–19.

3. Harms A, Brodersen DE, Mitarai N, Gerdes K. 2018. Toxins, Targets, and Triggers: An Overview of Toxin-Antitoxin Biology. Molecular Cell 70:768–784.

4. Page R, Peti W. 2016. Toxin-antitoxin systems in bacterial growth arrest and persistence. Nature Chemical Biology 12:208–214.

5. Schuster CF, Bertram R. 2013. Toxin-antitoxin systems are ubiquitous and versatile modulators of prokaryotic cell fate. FEMS Microbiol Lett 340:73–85.

6. Yamaguchi Y, Park J-H, Inouye M. 2011. Toxin-Antitoxin Systems in Bacteria and Archaea. Annual Review of Genetics 45:61–79.

7. Gerdes K, Christensen SK, Lobner-Olesen A. 2005. Prokaryotic toxin-antitoxin stress response loci. Nat Rev Microbiol 3:371–82.

8. Winther KS, Gerdes K. 2012. Regulation of enteric vapBC transcription: induction by VapC toxin dimer-breaking. Nucleic Acids Res 40:4347–57.

9. Jurėnas D, Van Melderen L. 2020. The Variety in the Common Theme of Translation Inhibition by Type II Toxin–Antitoxin Systems. Frontiers in Genetics 11.

10. Jiang Y, Pogliano J, Helinski DR, Konieczny I. 2002. ParE toxin encoded by the broad-host-range plasmid RK2 is an inhibitor of Escherichia coli gyrase. Mol Microbiol 44:971–9.

11. Hallez R, Geeraerts D, Sterckx Y, Mine N, Loris R, Van Melderen L. 2010. New toxins homologous to ParE belonging to three-component toxin–antitoxin systems in Escherichia coli O157:H7. Molecular Microbiology 76:719–732.

12. Christensen-Dalsgaard M, Gerdes K. 2006. Two higBA loci in the Vibrio cholerae superintegron encode mRNA cleaving enzymes and can stabilize plasmids. Mol Microbiol 62:397–411.

13. Lima-Mendez G, Oliveira Alvarenga D, Ross K, Hallet B, Van Melderen L, Varani AM, Chandler M. 2020. Toxin-Antitoxin Gene Pairs Found in Tn3 Family Transposons Appear To Be an Integral Part of the Transposition Module. mBio 11:e00452–20.

14. Dy RL, Przybilski R, Semeijn K, Salmond GPC, Fineran PC. 2014. A widespread bacteriophage abortive infection system functions through a Type IV toxin–antitoxin mechanism. Nucleic Acids Research 42:4590–4605.

15. Fineran PC, Blower TR, Foulds IJ, Humphreys DP, Lilley KS, Salmond GP. 2009. The phage abortive infection system, ToxIN, functions as a protein-RNA toxin-antitoxin pair. Proc Natl Acad Sci U S A 106:894–9.

16. Harms A, Fino C, Sørensen MA, Semsey S, Gerdes K. 2017. Prophages and Growth Dynamics Confound Experimental Results with Antibiotic-Tolerant Persister Cells. mBio 8:e01964–17.

17. Wang X, Kim Y, Hong SH, Ma Q, Brown BL, Pu M, Tarone AM, Benedik MJ, Peti W, Page R, Wood TK. 2011. Antitoxin MqsA helps mediate the bacterial general stress response. Nat Chem Biol 7:359–66.

18. Fraikin N, Rousseau CJ, Goeders N, Van Melderen L. 2019. Reassessing the Role of the Type II MqsRA Toxin-Antitoxin System in Stress Response and Biofilm Formation: mqsA Is Transcriptionally Uncoupled from mqsR. mBio 10:e02678–19.

19. LeRoux M, Culviner PH, Liu YJ, Littlehale ML, Laub MT. 2020. Stress Can Induce Transcription of Toxin-Antitoxin Systems without Activating Toxin. Molecular Cell doi:https://doi.org/10.1016/j.molcel.2020.05.028.

20. Dorr T, Vulic M, Lewis K. 2010. Ciprofloxacin causes persister formation by inducing the TisB toxin in Escherichia coli. PLoS Biol 8:e1000317.

21. Pontes MH, Groisman EA. 2020. A Physiological Basis for Nonheritable Antibiotic Resistance. mBio 11:e00817–20.

22. Christensen-Dalsgaard M, Jorgensen MG, Gerdes K. 2010. Three new RelE-homologous mRNA interferases of Escherichia coli differentially induced by environmental stresses. Mol Microbiol 75:333–48.

23. Kirkpatrick CL, Martins D, Redder P, Frandi A, Mignolet J, Chapalay JB, Chambon M, Turcatti G, Viollier PH. 2016. Growth control switch by a DNA-damage-inducible toxin–antitoxin system in Caulobacter crescentus. Nature Microbiology 1:16008.

24. Lasker K, Mann TH, Shapiro L. 2016. An intracellular compass spatially coordinates cell cycle modules in Caulobacter crescentus. Curr Opin Microbiol 33:131–139.

25. Tsokos CG, Laub MT. 2012. Polarity and cell fate asymmetry in Caulobacter crescentus. Curr Opin Microbiol 15:744–50.

26. Fumeaux C, Radhakrishnan SK, Ardissone S, Theraulaz L, Frandi A, Martins D, Nesper J, Abel S, Jenal U, Viollier PH. 2014. Cell cycle transition from S-phase to G1 in Caulobacter is mediated by ancestral virulence regulators. Nat Commun 5:4081.

27. Gora KG, Tsokos CG, Chen YE, Srinivasan BS, Perchuk BS, Laub MT. 2010. A cell-type-specific protein-protein interaction modulates transcriptional activity of a master regulator in Caulobacter crescentus. Mol Cell 39:455–67.

28. Tan MH, Kozdon JB, Shen X, Shapiro L, McAdams HH. 2010. An essential transcription factor, SciP, enhances robustness of Caulobacter cell cycle regulation. Proceedings of the National Academy of Sciences 107:18985.

29. Ryan KR, Huntwork S, Shapiro L. 2004. Recruitment of a cytoplasmic response regulator to the cell pole is linked to its cell cycle-regulated proteolysis. Proc Natl Acad Sci U S A 101:7415–20.

30. da Rocha RP, Paquola AC, Marques Mdo V, Menck CF, Galhardo RS. 2008. Characterization of the SOS regulon of Caulobacter crescentus. J Bacteriol 190:1209–18.

31. Dufour D, Mankovskaia A, Chan Y, Motavaze K, Gong S-G, Lévesque CM. 2018. A tripartite toxin-antitoxin module induced by quorum sensing is associated with the persistence phenotype in Streptococcus mutans. Molecular Oral Microbiology 33:420–429.

32. Vang Nielsen S, Turnbull KJ, Roghanian M, Bærentsen R, Semanjski M, Brodersen DE, Macek B, Gerdes K. 2019. Serine-Threonine Kinases Encoded by Split hipA Homologs Inhibit Tryptophanyl-tRNA Synthetase. mBio 10:e01138–19.

33. Bordes P, Cirinesi A-M, Ummels R, Sala A, Sakr S, Bitter W, Genevaux P. 2011. SecB-like chaperone controls a toxin–antitoxin stress-responsive system in *Mycobacterium tuberculosis*. Proceedings of the National Academy of Sciences 108:8438–8443.

34. Simon R, Priefer U, Pühler A. 1983. A broad host range mobilization system for *in vivo* genetic engineering: transposon mutagenesis in Gram negative bacteria. Bio/Technology 1:784–791.

35. Miller JH. 1972. Assay of β-galactosidase, p 352–355. In Miller JH (ed), Experiments in Molecular Genetics. Cold Spring Harbor Laboratory Press.

36. Radhakrishnan SK, Thanbichler M, Viollier PH. 2008. The dynamic interplay between a cell fate determinant and a lysozyme homolog drives the asymmetric division cycle of Caulobacter crescentus. Genes Dev 22:212–25.

37. Solovyev V, Salamov A. 2011. Automatic Annotation of Microbial Genomes and Metagenomic Sequences., p 61–78, Metagenomics and its Applications in Agriculture, Biomedicine and Environmental Studies. Nova Science Publishers.

38. Gautheret D, Lambert A. 2001. Direct RNA motif definition and identification from multiple sequence alignments using secondary structure profiles11Edited by J. Doudna. Journal of Molecular Biology 313:1003–1011.

39. Macke TJ, Ecker DJ, Gutell RR, Gautheret D, Case DA, Sampath R. 2001. RNAMotif, an RNA secondary structure definition and search algorithm. Nucleic Acids Research 29:4724–4735.

40. Nikolskaya AN, Galperin MY. 2002. A novel type of conserved DNA-binding domain in the transcriptional regulators of the AlgR/AgrA/LytR family. Nucleic Acids Res 30:2453–9.

41. Cserzo M, Wallin E, Simon I, von Heijne G, Elofsson A. 1997. Prediction of transmembrane alpha-helices in prokaryotic membrane proteins: the dense alignment surface method. Protein Eng 10:673–6.

42. Almagro Armenteros JJ, Tsirigos KD, Sønderby CK, Petersen TN, Winther O, Brunak S, von Heijne G, Nielsen H. 2019. SignalP 5.0 improves signal peptide predictions using deep neural networks. Nature Biotechnology 37:420–423.

43. Maddocks SE, Oyston PCF. 2008. Structure and function of the LysR-type transcriptional regulator (LTTR) family proteins. Microbiology 154:3609–3623.

44. Huang CY, Gonzalez-Lopez C, Henry C, Mijakovic I, Ryan KR. 2020. hipBA toxin-antitoxin systems mediate persistence in Caulobacter crescentus. Scientific Reports 10:2865.

45. Domian IJ, Quon KC, Shapiro L. 1997. Cell type-specific phosphorylation and proteolysis of a transcriptional regulator controls the G1-to-S transition in a bacterial cell cycle. Cell 90:415–24.

46. Modell JW, Hopkins AC, Laub MT. 2011. A DNA damage checkpoint in Caulobacter crescentus inhibits cell division through a direct interaction with FtsW. Genes Dev 25:1328–43.

47. Modell JW, Kambara TK, Perchuk BS, Laub MT. 2014. A DNA damage-induced, SOS-independent checkpoint regulates cell division in Caulobacter crescentus. PLoS Biol 12:e1001977.

48. Zhou B, Schrader JM, Kalogeraki VS, Abeliuk E, Dinh CB, Pham JQ, Cui ZZ, Dill DL, McAdams HH, Shapiro L. 2015. The global regulatory architecture of transcription during the Caulobacter cell cycle. PLoS Genet 11:e1004831.

49. Laub MT, Chen SL, Shapiro L, McAdams HH. 2002. Genes directly controlled by CtrA, a master regulator of the Caulobacter cell cycle. Proc Natl Acad Sci U S A 99:4632–7.

50. Thanbichler M, Iniesta AA, Shapiro L. 2007. A comprehensive set of plasmids for vanillate- and xylose-inducible gene expression in Caulobacter crescentus. Nucleic Acids Res 35:e137.

51. Evinger M, Agabian N. 1977. Envelope-associated nucleoid from Caulobacter crescentus stalked and swarmer cells. J Bacteriol 132:294–301.

